# Examination of low-intensity focused ultrasound parameters for human neuromodulation

**DOI:** 10.1101/2024.09.23.614460

**Authors:** Areej Ennasr, Gabriel Isaac, Andrew Strohman, Wynn Legon

**Author notes:** Correspondence: Wynn Legon,; 1 Riverside Circle Roanoke, VA, 24016, USA. denotes co-1^st^ authorship (Areej Ennasr, Gabriel Isaac).

## Abstract

**Background:** Low-intensity focused ultrasound (LIFU) is a promising form of non-invasive neuromodulation characterized by a rich parameter space that includes intensity, duration, duty cycle and pulsing strategy. The effect and interaction of these parameters to affect human brain activity is poorly understood. A better understanding of how parameters interact is critical to advance LIFU as a potential therapeutic.

**Objective/Hypothesis:** To determine how different parameters, including intensity, duration, and duty cycle interact to produce neuromodulation effects in the human brain. Further, this study assesses the effect of pulsing versus continuous ultrasound. We hypothesize that higher duty cycles will confer excitation. Increasing intensity or duration will increase the magnitude of effect. Pulsing LIFU will not be more effective than continuous wave ultrasound.

**Methods:** N = 18 healthy human volunteers underwent 20 different parameter combinations that included a fully parametrized set of two intensities (I_SPPA_: 6 & 24 W/cm^2^), five duty cycles (1, 10, 30, 50, 70%) and two durations (100, 500 msec) with a constant pulse repetition frequency of 1 kHz delivered concurrently with transcranial magnetic stimulation (TMS) to the primary motor cortex (M1). Four of these parameter combinations were also delivered continuously, matched on the number of cycles. Motor-evoked potential (MEP) amplitude was the primary outcome measure. All parameter combinations were collected time-locked to MEP generation.

**Results:** There was no evidence of excitation from any parameter combination. 3 of the 24 parameter sets resulted in inhibition. The parameter set that resulted in the greatest inhibition (∼ 30%) was an intensity of 6W/cm^2^ with a duty cycle of 30% and a duration of 500 msec. A three-way ANOVA revealed an interaction of intensity and duty cycle. The analysis of continuous versus pulsed ultrasound revealed a 3-way interaction of intensity, pulsing, and the number of cycles such that under the 6W/cm^2^ condition higher cycles of pulsed ultrasound resulted in inhibition whereas lower number of cycles using continuous LIFU resulted in inhibition.

**Conclusions:** LIFU to M1 in humans, in the range employed, either conferred inhibition or had no effect. Significant excitation was not observed. In general, lower intensity looks to be more efficacious for inhibition that depends on duration. Finally, pulsed ultrasound looks to be more effective for inhibition as compared to continuous wave after controlling for total energy delivered.

## INTRODUCTION

Low-intensity focused ultrasound (LIFU) is a non-invasive neuromodulatory approach that uses mechanical energy to reversibly modulate neuronal activity with high spatial resolution and adjustable depth of focus [1,2]. LIFU has been extensively studied in small [3–5] and large animals [6–9] including humans [10–13]. It has been demonstrated to produce safe [14,15] and reversible inhibition and excitation [8,16] in different parts of the cortex [12,13,17–20] and sub-cortical structures [1,21] including emerging clinical applications [22–26]. Despite these results it remains unclear how and if different parameters affect neuromodulation. LIFU has a rich parameter space that includes amplitude (pressure) or intensity (the square of amplitude), duration, duty cycle, and pulse repetition frequency. Several of these parameters have been empirically tested in slice models [27], single-cell [28], invertebrate [29], small animal [3–5,30], and large animal [31] and there are now a few studies in human [13,32,33]. Furthermore, theoretical models have also been developed to help explain results found in small animal [34]. In humans, Fomenko et al. (2020) found that 10% duty cycle conferred the greatest inhibition and that longer sonication durations were more effective for inhibition while changes in pulse repetition frequency had no effect of the amplitude of motor-evoked potentials (MEPs) when duty cycle was controlled for [13]. Despite these important results, this study did not examine how and if any of these parameters interact to affect results as only one parameter was adjusted at a time. The small animal work and theoretical modelling propose that parameters may interact to affect results but this has never been empirically tested in humans. This is critical to determine if results in small animal work translate to humans and also for direct human evidence that will ultimately help with translation to potential clinical therapeutic applications. Here, we examine a full parametric model of 500 kHz LIFU directed at the primary motor cortex (M1) in humans for effect on the amplitude of the transcranial magnetic stimulation (TMS) derived MEP [10]. We examined two different spatial peak pulse average intensities (I_SPPA_: 6, 24 W/cm^2^), five duty cycles (1, 10, 30, 50, 70 %) and two durations (100, 500 msec) at a constant pulse repetition frequency of 1kHz. In addition, we examined how and if pulsing LIFU from two different parameters combinations compares to delivering them as a continuous wave while keeping the total number of cycles constant across conditions. The effect of pulsing ultrasound has been studied but results are equivocal [4,35], though this has not yet been examined in humans. We hypothesized that lower duty cycles would confer inhibition and higher duty cycles would confer excitation based upon small animal work and theoretical models [34]. We further hypothesized that increasing total energy through either an increase in intensity or duration would increase any effect (inhibition or excitation) from the different duty cycles tested. Finally, we hypothesized no difference between pulsed versus continuous ultrasound under the hypothesis that total amount of energy is the determining factor for neuromodulation and not its method of delivery.

## MATERIALS & METHODS

### Participants

The Institutional Review Board at Virginia Tech approved all experimental procedures (IRB #21-882). Participants had to meet all inclusion/exclusion criteria provided written informed consent to all aspects of the study. Inclusion criteria were males and females ages 18-65 who were physically and neurologically healthy with no history of neurological disorder. Exclusion criteria included contraindications to imaging (magnetic resonance imaging (MRI) & computed tomography (CT)), a history of neurologic disorder or head injury resulting in loss of consciousness for >10 minutes, any drug dependence, and any active medical disorder or current treatment with potential central nervous system effects.

### Overall Study Design

The overall design consisted of up to five separate visits on five separate days. The first visit consisted of anatomical MRI and CT scans for acoustic modelling and neuronavigation purposes (details below) in addition to baseline questionnaires. The subsequent visits were formal testing sessions of the LIFU parameter combinations. For the LIFU testing sessions, subjects received up to 24 unique parameter combinations. There was a minimum of 24 hours between each session. The number of parameter combinations completed per session varied between subjects based upon experimental and time constraints. Additionally, report of symptoms questionnaires were collected after each visit. Each LIFU testing session lasted roughly 2.5 hours. A final report of symptoms questionnaire was collected via email one month following final study participation. The overall design is depicted in **Figure 1A**.

**Figure 1.**
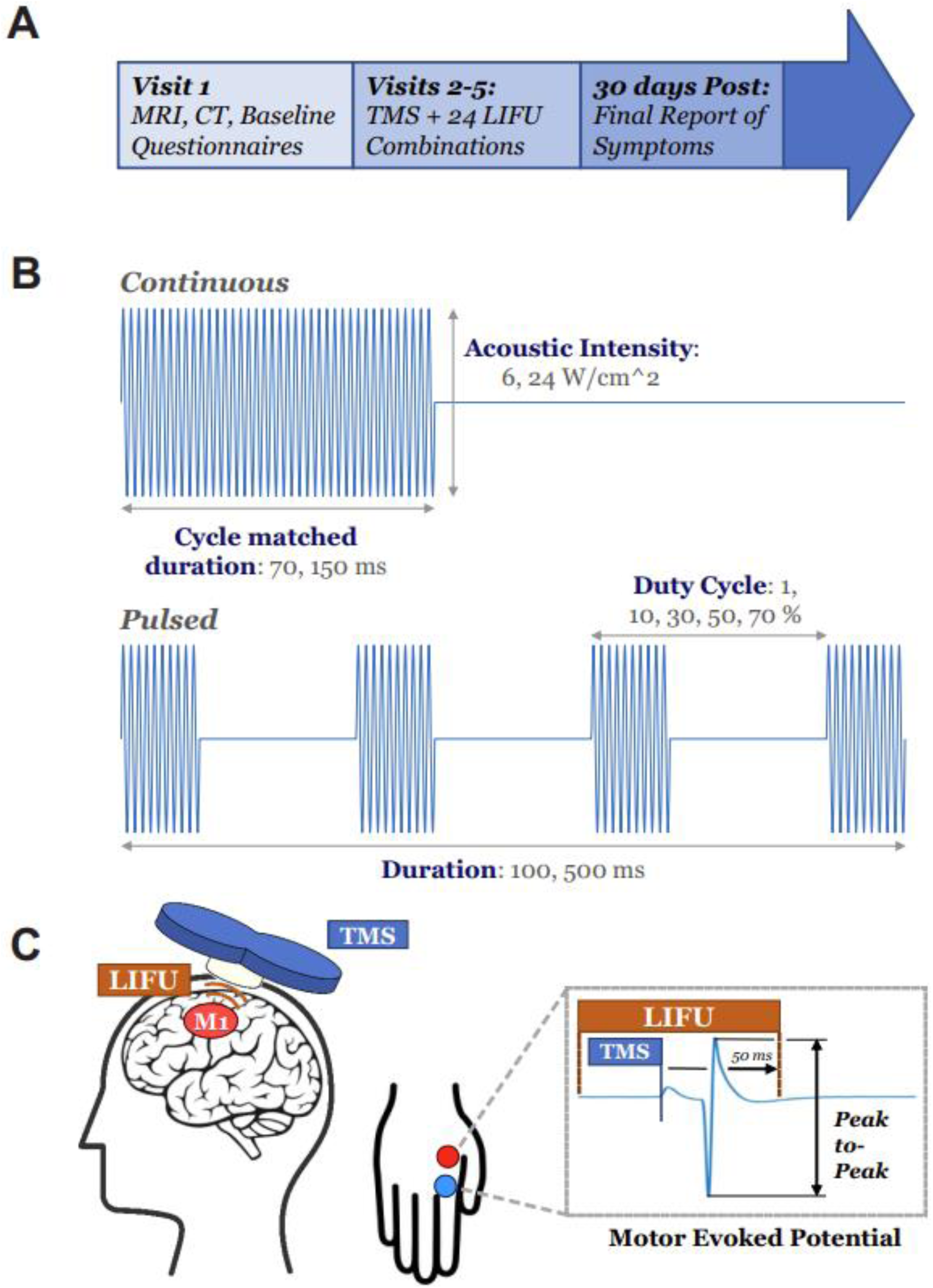
Experimental design. **(A)** Timeline summarizing the overall experimental design. During the first visit, participants underwent structural MRI and CT scans and complete baseline questionnaires. Over subsequent visits (up to four), 24 unique combinations of LIFU were tested. Follow-up symptom questionnaires were sent 30 days after final visit. **(B)** Schematic depicting the LIFU parameters tested: duration (100, 500 msec), duty cycle (1, 10, 30, 50, 70 %), acoustic intensity (I_SPPA_: 6, 24 W/cm^2^), and cycle matched continuous-wave duration (70, 150ms), at a constant pulse repetition frequency of 1 kHz. **(C)** Schematic illustrating simultaneous LIFU and TMS stimulation to the M1, inducing a motor evoked potential (MEP) recorded at the first dorsal interosseous muscle. The TMS pulse was time-locked to 50 msec before the end of LIFU stimulation for all parameter sets tested and MEP amplitudes were used to examine changes in corticospinal excitability.

### MRI and CT Acquisition

Magnetic resonance imaging (MRI) and computed tomography (CT) scans were collected for neuronavigation and acoustic modeling to accurately estimate beam volume and energy delivery to the target region of the M1. MRI data were acquired on a Siemens 3T Prisma scanner (Siemens Medical Solutions, Erlangen, Germany) at the Fralin Biomedical Research Institute’s Human Neuroimaging Laboratory. Anatomical scans were acquired using a T1-weighted MPRAGE sequence with a TR = 1400 ms, TI = 600 ms, TE = 2.66 ms, flip angle = 12 (degrees), voxel size = 0.5×0.5×1.0 mm, FoV read = 245 mm, FoV phase of 87.5%, 192 slices, ascending acquisition. CT scans were collected with a Kernel = Hr60 in the bone window, FoV = 250 mm, kilovolts (kV) = 120, rotation time = 1 second, delay = 2 seconds, pitch = 0.55, caudocranial image acquisition order, 1.0 mm image increments for a total of 121 images and scan time of 13.14 seconds.

### Parameters

The parameters tested included two different spatial peak pulse average intensities (I_SPPA_: 6, 24 W/cm^2^), five duty cycles (1, 10, 30, 50, 70 %) and two durations (100, 500 msec), at a constant pulse repetition frequency of 1kHz, resulting in a total of 20 pulsed LIFU conditions. Two continuous-wave (CW) durations (70 msec and 150 msec) were also tested at 6 and 24 W/cm^2^ I_SPPA_, resulting in a total of 4 CW conditions. Notably, these CW durations were cycle-matched to the pulsed LIFU combinations of 70% duty cycle for 100 msec (pulsed conditions 5, 15) and 30% duty cycle for 500 msec (pulsed conditions 8, 18), respectively, to allow for comparison between pulsing schemes. This resulted in a total of 24 LIFU conditions tested. See **Figure 1B**.

### Motor evoked potential collection

#### Hotspot targeting and Baseline MEP collection

The target location for stimulation was initially set to the “hand knob” region of each participant’s left pre-central gyrus to find the ‘hot spot’ for the first dorsal interosseous (FDI) muscle using an MR-guided neuronavigation system (Brainsight, Rogue Research, Montreal, QUE, CAN). TMS stimulation was initially set to a stimulator output of 45%, increasing by 3% increments as needed to achieve at least 3 out of 5 MEP peak-to-peak amplitudes of at least 50µV. Additional locations 1 cm anterior, posterior, lateral, and medial to the initial location were also tested using the same criteria. Once the hotspot was determined (without the LIFU transducer), the TMS stimulation was re-thresholded with the LIFU transducer fixed underneath the TMS coil (see Legon et al. 2018 [10] for the effect of the transducer on the TMS generated electric field in the brain), increasing the stimulator output until an MEP amplitude of 750 mV was attained. If an MEP amplitude of 750 µV could not be achieved with a stimulator output of 100%, the visit was terminated and the subject was not included in the study. 750 mV was chosen as the baseline amplitude so that decrease and increase of MEP amplitude could be readily detected. After thresholding, pre-LIFU MEP amplitudes were collected using 20 TMS stimulations at an interstimulus interval (ISI) of 10 seconds. The average of these 20 MEPs served as the baseline for each participant and used to normalize and compare the effects of each unique LIFU parameter set. All 20 MEPs regardless of amplitude were used in this average. Once baseline measures were obtained, the visit proceeded with LIFU parameter testing. A new baseline was collected at the beginning of each visit to account for potential day-to-day variability in motor cortical excitability.

#### Electromyography (EMG)

Surface electromyography (EMG) data was collected by placing adhesive electrodes (Medi-Trace® 530 series) in a belly-tendon montage over the first dorsal interosseous (FDI) muscle of the right hand, with ground placed on the olecranon process of the ipsilateral ulna. All electrodes were secured with tape. EMG data was continuously recorded using a DC amplifier (BrainSight, Rogue Research, Montreal, QUE, CAN) sampled at 1 kHz and stored on a PC for later offline analysis. EMG analysis was performed using custom-made scripts written in MATLAB (Mathworks Natick MA USA).

#### Concurrent LIFU & TMS

All MEPs were generated using a commercial figure eight TMS coil of two coplanar coil windings of equal size (D70^2^ alpha coil, Magstim Inc., UK). A custom single-element LIFU transducer was mounted at the center of the coil intersection using a custom 3D-printed fixture [10]. The TMS/LIFU apparatus placement was tracked using an MR-guided neuronavigation system (BrainSight, Rogue Research, Montreal, QUE, CAN). During the concurrent stimulation, ultrasound transmission gel (Aquasonic 100; Parker Laboratories Inc., NJ, USA) was applied between the transducer and the parted hair/skin of the participant’s head to aid energy transfer to the brain. The TMS coil was positioned at a 45-degree angle from the midline, with the coil handle facing posteriorly producing a posterior to anterior current in the brain. For all parameter combinations, TMS was time-locked to 50 msec before the end of the LIFU stimulation to ensure LIFU overlapped with the timing of the generation of the MEP (**Figure 1C**). A total of 20 LIFU/TMS stimulations were delivered for each parameter combination with an ISI of 10 seconds.

The order of the 24 parameters was randomized for each participant prior to data collection visits to help account for potential carry-over-effects between conditions. Parameter combinations were tested in sequence with no rest period between tests unless no-go criteria were met. Go/no-go criteria were put in place to help mitigate potential carry-over effects between the parameter sets. The go/no-go criteria were as follows: If the MEP amplitudes from the previous parameter set consistently (3 out of the last 5 MEPs) fell below 100 µV, a 10-minute wait was implemented before proceeding to the next combination. After 10 minutes, 20 additional TMS stimulations without LIFU at the original final stimulator output were conducted to confirm that the MEP amplitudes had been restored to baseline magnitudes. If the MEP amplitudes were not restored, another 10-minute wait was implemented, and the process was repeated until the MEPs were restored or until the allotted time for the visit was exhausted (The subjects, sessions and conditions where go/no-go were used are provided in **Supplemental Table 2**).

#### LIFU Waveform and Delivery

A custom-made single-element focused transducer with a center frequency of 500 kHz, a 30 mm diameter, and a 31 mm focal length was used (**Figure 2 A&B**). The LIFU waveforms were generated using a two-channel 2-MHz function generator (4078B, B&K Precision Instruments, CA, USA). Channel 1 was configured to generate a 5Vp-p square wave frequency of 1 kHz waveform in burst mode triggered externally. The wave parameters of duty cycle, duration, and intensity were adjusted externally using a custom MATLAB script and were used to gate channel 2 which was set to generate a 500 kHz sine wave. Based on transducer characterization, the amplitude for channel 2 was set to either 0.21 Volts peak-to-peak (Vpp) or 0.44 Vpp, which corresponds to intensity parameters of 6 W/cm² or 24 W/cm² (I_SPPA_), respectively (as determined by empirical acoustic testing in free water). For testing continuous parameter settings only channel 1 was used with a frequency of 500 kHz, an amplitude of either 0.21 Vpp or 0.44 Vpp, and the number of cycles set to 35,000 or 75,000. The number of cycles was determined by matching the total energy delivered between the selected continuous and pulsed conditions. This was based on the total “on-time” for the pulsed conditions of 100msec at 70% DC and 500msec at 30% DC, respectively. For both the pulsed and continuous conditions, the output of channel 2 was sent through a 100 W linear RF amplifier (E&I 2100L; Electronics & Innovation) before being sent to the transducer through its corresponding matching network.

**Figure 2.**
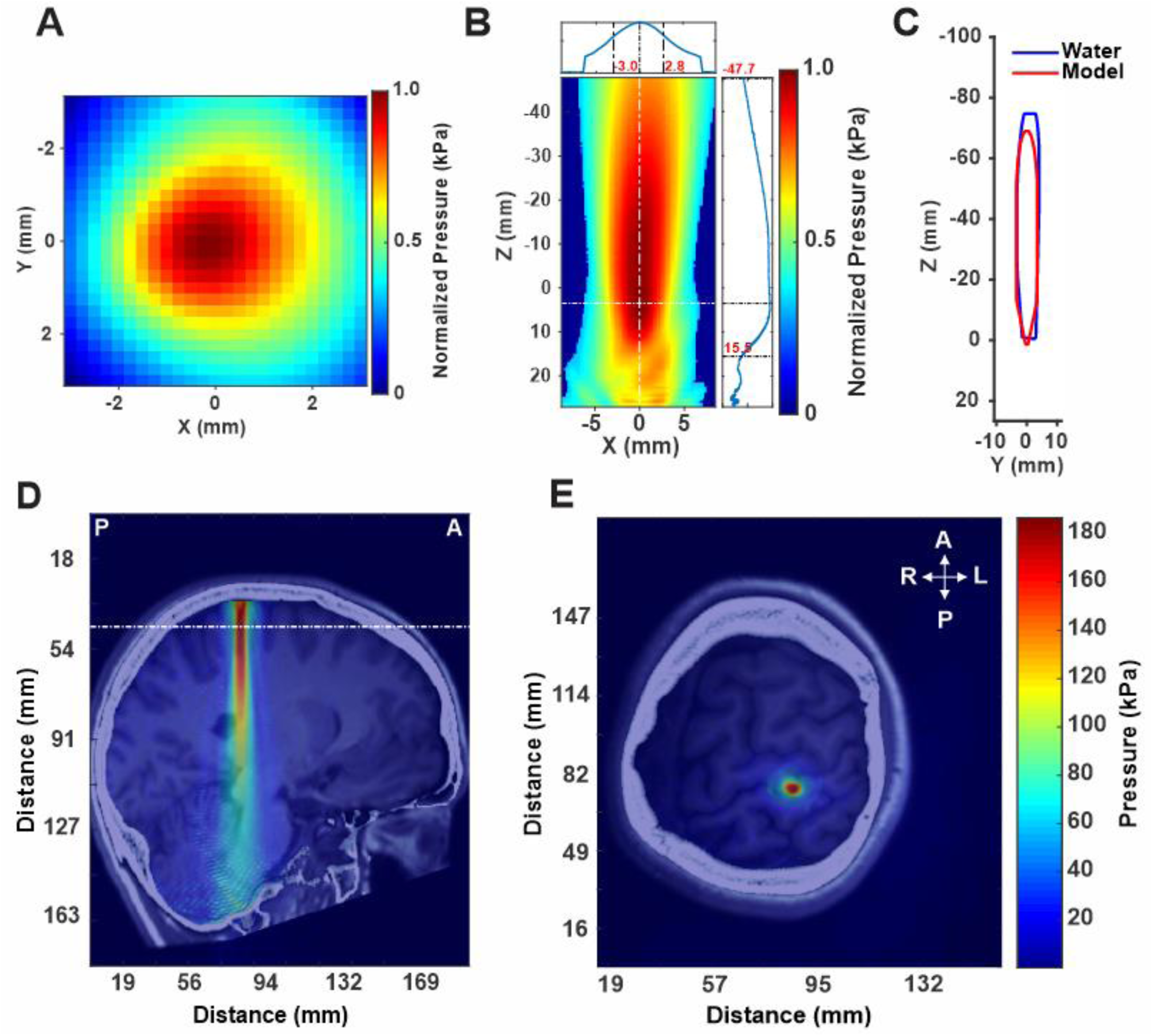
Transducer Characteristics and Modelling. **A.** Pseudo-color map of free water XY lateral beam pressure at Z maximum of the 500 kHz single-element transducer **B.** Pseudo-color map of free water XZ axial beam pressure. The numbers in red represent the limits of the full-width half-maximum (FWHM) pressure in millimeters from the focal spot. The transducer face is at Z = 31 millimeters (mm). **C.** FWHM overlay of the empirically measured XZ axial beam in acoustic test tank (blue) and the modeled beam (red) used for acoustic modeling. **D, E.** Acoustic modelling results using MRI and CT scans from the same representative subject. Warmer colors indicate more pressure (kPa). Images depict the sagittal (**D**) and transverse (**E**) views. The transverse cross-section in E is taken from the white line in the sagittal view. R = right, L = left, P = posterior, A = anterior.

#### Empirical Acoustic Field Mapping

The acoustic intensity of the ultrasound waveform was measured using an acoustic test tank filled with deionized, degassed, and filtered water (Precision Acoustics Ltd.). A calibrated hydrophone (HNR-0500, Onda Corp.) was fixed on a motorized stage to measure the pressure generated by the ultrasound transducer. The transducer was positioned using a custom setup and aligned to ensure the hydrophone was perpendicular to its surface. Planar scans in the XY and YZ planes were conducted with an isotropic resolution of 0.25 × 0.25 mm. Additionally, a voltage sweep ranging from 20 to 250 mVpp in 10 mVpp increments was performed to determine the input voltage required to achieve the desired extracranial pressure.

### Modelling

#### Acoustic Modelling

Computational models were developed for both intensity conditions (6, 24 W/cm^2^ I_SPPA_) using individual subject MR and CT images to evaluate the wave propagation of LIFU across the skull and the resultant intracranial acoustic pressure maps. Simulations were performed using the k-Wave MATLAB toolbox [36], which uses a pseudospectral time domain method to solve discretized wave equations on a spatial grid. CT images were used to construct the acoustic model of the skull, while MR images were used to target LIFU at M1, based on individual brain anatomy. Details of the modeling parameters can be found in Legon et al. (2018) [1]. Briefly, CT and MR images were first co-registered and then up-sampled for acoustic simulations. The skull was extracted manually using a threshold intensity value and the intracranial space was assumed to be homogenous as ultrasound reflections between soft tissues are small [37]. Acoustic parameters were calculated from CT data assuming a linear relationship between skull porosity and the acoustic parameters [38,39]. The computational model of the ultrasound transducer used in simulations was constructed to recreate empirical acoustic pressure maps of focused ultrasound transmitted in the acoustic test tank similar to previous work [1,10,40,41] (**Figure 2C**).

#### Thermal Modelling

Thermal models were generated using the modified mixed-domain method (msecOUND)1 with a 3-layer skull model (cortical-trabecular-cortical) from Benchmark 6 in Aubry et al., (2022) to solve for the temperature fields based on the acoustic field and the bioheat equation using k-wave Diffusion3 [42]. We performed a thermal simulation using the parameter set with the highest duty cycle, longest duration, and highest intensity (70% duty cycle (DC), 500msec pulse duration, I_SPPA_ = 24 W/cm^2^ or 888 kilopascals) at a 10 Hz pulse repetition frequency (PRF) and a 10 second interstimulus interval (ISI) for 20 stimulations. This thermal simulation assumes a 65% energy loss through the skull where 100% of the energy loss is absorption (not scattering) and is thus converted to heating, thus providing a conservative estimate of thermal rise. The simulation was run at 10 Hz PRF as opposed to 1 kHz PRF as the higher PRF was too computationally demanding to fully simulate with current resources. Using longer pulse durations have been shown to minimally alter modelling outcomes when total energy delivery is held constant [43].

#### Statistical Analysis

MEP amplitudes were measured as the peak-to-peak response. Assumptions were evaluated using Bartlett’s test for sphericity with a threshold of p < 0.05. Variables with significant results were tested using the appropriate non-parametric test. For each subject, MEP amplitude of the 20 stimulations for each trial was averaged. In order to test this, we ran a paired t-test across 2 and 3 to evaluate if there were differences between baseline MEP amplitudes at a threshold of p < 0.05. To determine if any of the pulsed parameter sets were different from baseline, we performed separate paired t-tests comparing normalized baseline MEP values with normalized MEP for each parameter set at a Bonferroni correction level of p < 0.0025 (0.05/20). To test for significant differences between the components of each parameter, a 3-way repeated measures analysis of variance (ANOVA) was performed with main factors INTENSITY (I_SPPA_: 6 W/cm^2^, 24 W/cm^2^), DUTY CYCLE (1%, 10%, 30%, 50%, 70%), and DURATION (100 msec, 500 msec) at a threshold of p < 0.05. A separate 3-way repeated measures ANOVA was run to test for pulsed versus continuous parameter delivery with main factors INTENSITY (I_SPPA_: 6 W/cm^2^, 24 W/cm^2^), NUMBER OF CYCLES (35000, 75000), and PULSING (pulsed, continuous) at a threshold of p < 0.05. Interactions and main effects were examined using appropriate post-hoc testing.

## RESULTS

### Participants

Of the 23 people enrolled, one participant was lost to follow-up after the first session, and four participants failed to achieve an MEP of at least 750 µV at baseline, so 18 participants proceeded through the remainder of the visits. Of the 18 healthy participants (25.6 years ± 4.4 years; range (20-38); M/F 10/8), all 18 completed the 20 pulsed conditions, while 17 also completed the continuous LIFU parameter sets. The number of parameter combinations tested per visit ranged between 4 and 17 due to various experimental limitations. While the majority of participants were able to complete all parameter combinations over two visits, four participants required three visits, and two participants required four visits (see **Supplemental Table 1**). Causes for requiring extra sessions included greater time required to find the M1 target spot or overheating of the TMS coil, requiring longer breaks between LIFU/TMS delivery in order for the TMS coil to cool down. Additionally, 4 subjects met no-go criteria during at least one parameter set requiring a 10-minute wait period(s), extending the duration of their visits and thus limiting the total number of parameters sets that could be collected.

### LIFU Beam Characteristics

Empirical acoustic measurements were conducted to evaluate the transducer’s beam profile in free water. These measurements indicated a full-width half maximum (FWHM) lateral (XY) resolution of ± 2.9 mm at the axial (Z) beam focus (**Fig. 2A**). The FWHM axial (YZ) resolution was ± 31.6 mm, with a focal depth of 31 mm from the transducer’s exit plane (**Fig. 2B**). The comparison between the empirical axial FWHM measurements in free water and the modeled waveform showed good agreement, thereby validating the use of these measurements in acoustic models (**Fig. 2C**). An example of the ultrasound beam directed at M1 for one subject is shown in **Figure 2 D & E**.

### Estimated intracranial pressure

Two different intensities were delivered to M1. The *in vivo* mean ± SD pressure estimated at the M1 for the 6W/cm^2^ condition was 141.11 ± 66.74 kPa, with a range of 75.88 to 354.02 kPa. The *in vivo* mean ± SD pressure estimated at the M1 for the 24W/cm^2^ condition was 275.25 ± 130.19 kPa, ranging from 148.02 to 690.57 kPa. A paired t-test revealed a significant difference between the two intensity conditions (t(17) = 9.28, p < 0.0001) – validating that the two conditions were separable. The mean ± SD estimated skull attenuation was 69.02 ± 14.65% and ranged from 22.27 to 83.34%. The estimated individual and average intracranial pressures for both intensities are summarized in **Table 1**.

**Table 1.** Estimated intracranial pressure. Individual estimated LIFU transmission and intracranial pressure in the head M1 region, as well as mean ± SD across all participants.

| Participant # | Average Transmission (%) | Pressure inside head (kPa) |  |
| --- | --- | --- | --- |
|  |  | *Isppa* = 6 W/cm<sup>2</sup> | *Isppa* = 24 W/cm<sup>2</sup> |
| 1 | 33.68 | 153.39 | 299.2 |
| 2 | 40.52 | 184.54 | 359.98 |
| 3 | 22.94 | 104.48 | 203.81 |
| 4 | 16.66 | 75.88 | 148.02 |
| 5 | 32.86 | 149.65 | 291.92 |
| 6 | 23.52 | 107.12 | 208.96 |
| 7 | 32.81 | 149.44 | 291.5 |
| 8 | 31.37 | 142.89 | 278.73 |
| 9 | 24.62 | 112.15 | 218.76 |
| 10 | 24.93 | 113.55 | 221.49 |
| 11 | 25.07 | 114.18 | 222.73 |
| 12 | 23.39 | 106.52 | 207.78 |
| 13 | 26.03 | 118.57 | 231.28 |
| 14 | 49.58 | 225.83 | 440.51 |
| 15 | 35.46 | 161.51 | 315.04 |
| 16 | 17.65 | 80.38 | 156.79 |
| 17 | 77.73 | 354.02 | 690.57 |
| 18 | 21.54 | 98.08 | 191.33 |
| **Mean ± SD** | **30.98 ± 14.65** | **141.11 ± 66.74** | **275.25 ± 130.19** |

### Estimated Temperature Rise

Simulations applied the parameter set yielding the highest potential thermal load and thus a conservative estimate of heating using a 70% duty cycle (DC), 500 msec pulse duration, 24 W/cm² I_SPPA_ (888 kPa), and 10 Hz PRF, with a 10-second ISI for 20 stimulations. The results indicated that the peak temperature rise occurred in the trabecular bone of the skull, reaching 37.79°C, while the peak temperature in the brain tissue was 37.41° (**Figure 3 A & B**).

**Figure 3.**
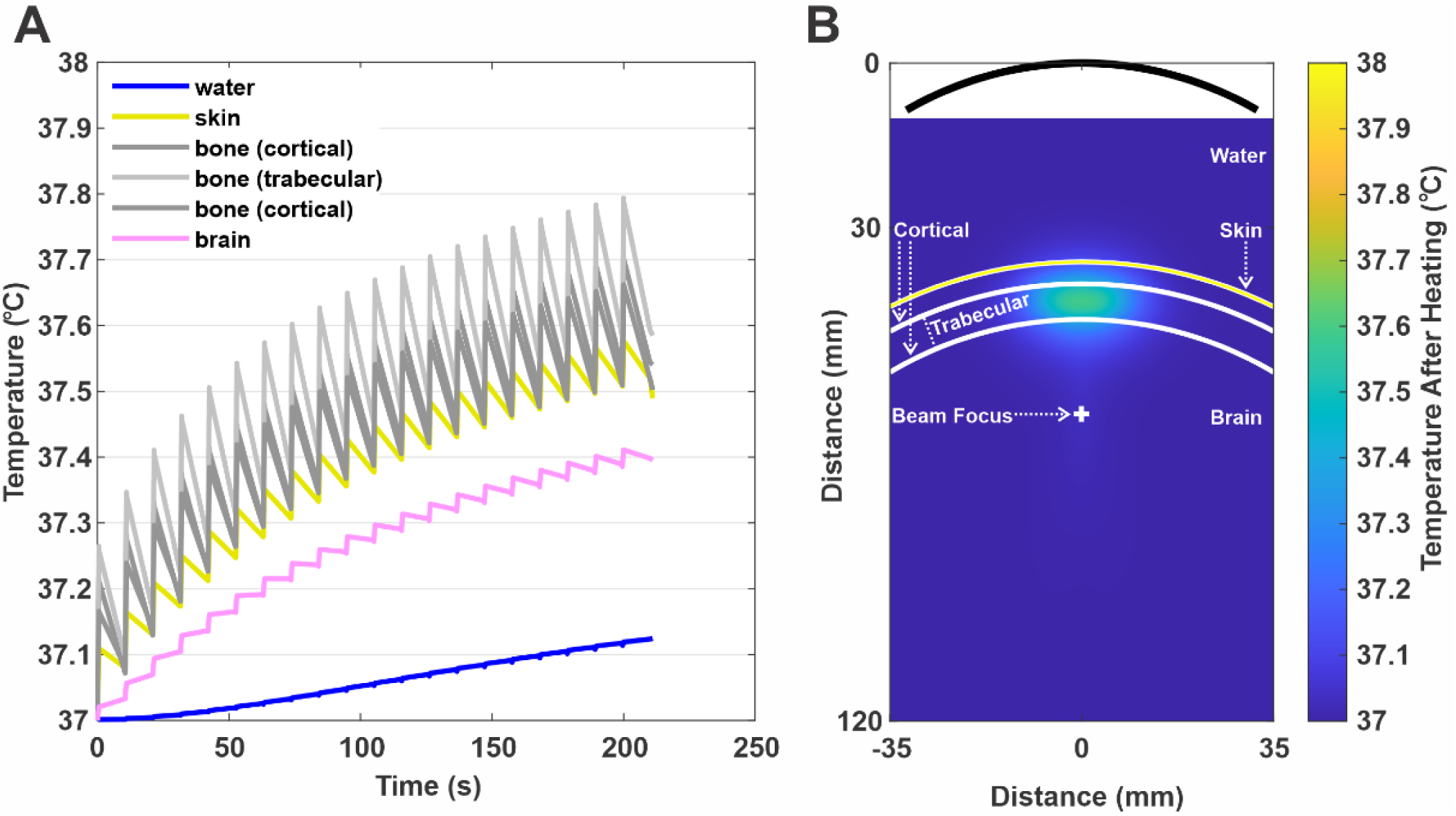
Thermal modeling. (**A**) Thermal rises over time in each of the 6 layers (water, skin, outer cortical bone, trabecular bone, inner cortical bone, brain) at the highest intensity (24 W/cm^2^ I_SPPA_ (888 kPa)), highest duration (500ms), highest duty cycle (70%) LIFU condition. Note the repeating spikes illustrating the 20 LIFU bursts spread across ∼200s. **B**) Pseudo-color map of simulated tissue heating at the final peak temperature (°C) illustrating spatial component of heating through the relevant 6 layers. The curved black line at the top of the figure represents the surface of the transducer.

### Parameter Results

The mean ± SD baseline MEP amplitudes for all participants across all visits were 625.7 ± 242.4 µV (see **Table 2**). We compared baseline MEP amplitude for session 2 and 3 as all participants were active in at least these sessions. The mean ± SD for session 2 and 3 were 574 ± 344 mV, 660 ± 376 mV, respectively. A paired samples t-test found no significant differences between session 2 and 3 baseline MEP amplitudes (t(17) = 1.11, p = 0.28). **Figure 4** shows average MEP amplitudes for baseline and 13 randomly tested LIFU conditions collected during one session in a single representative subject.

**Figure 4.**
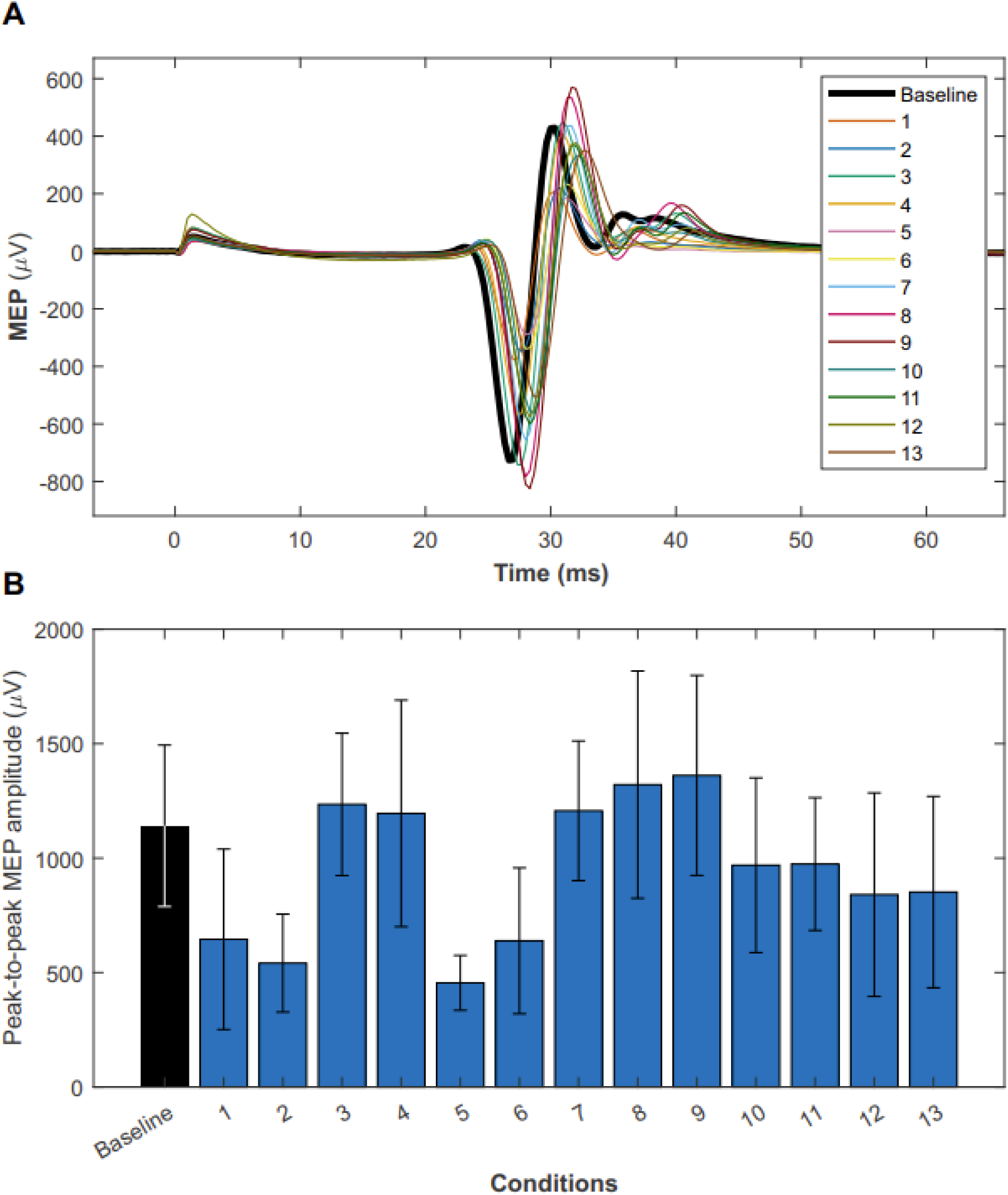
Example individual subject MEP amplitudes. Sample MEP data collection of a random 13 out of the 24 total parameter conditions in a single participant session. **(A)** EMG traces for each of the 13 parameter combinations are shown alongside baseline measurements, illustrating the ultrasonic modulation of MEP amplitudes. The TMS pulse is synchronized with time 0, and the MEP occurs approximately 30ms after this pulse. **(B)** The bar graph summarizes the mean ± SD MEP amplitudes by condition for the same set of LIFU parameters. The graph shows that some conditions appear to induce inhibition of the MEP, while others facilitate MEPs in the same subject. (Note: the parameters are arranged in the order they were tested during the session and do not reflect the parameter number from the group conditions).

**Table 2.** Participant baseline MEPs. Baseline average MEP (n=20) using TMS alone for each participant in every session before LIFU application.

| Participant # | Baseline MEP ± SD (mV) | | | |
| --- | --- | --- | --- | --- |
|  | Session 2 | Session 3 | Session 4 | Session 5 |
| 1 | 224±52 | 308±104 | - | - |
| 2 | 529±255 | 1008±566 | - | - |
| 3 | 550±375 | 392±284 | 782±550 | - |
| 4 | 1141±352 | 713±157 | - | - |
| 5 | 564±176 | 697±161 | - | - |
| 6 | 828±130 | 922±386 | 1291±284 | - |
| 7 | 958±349 | 560±137 | - | - |
| 8 | 178±97 | 400±220 | - | - |
| 9 | 184±38 | 292±291 | 265±210 | - |
| 10 | 1376±580 | 1704±716 | - | - |
| 11 | 519±280 | 315±138 | 458±169 | - |
| 12 | 528±151 | 879±410 | - | - |
| 13 | 168±101 | 1134±453 | - | - |
| 14 | 545±90 | 536±177 | - | - |
| 15 | 654±221 | 849±213 | - | - |
| 16 | 270±173 | 259±217 | 262±184 | - |
| 17 | 814±98 | 579±198 | 717±343 | 916±110 |
| 18 | 300±111 | 336±115 | - | - |
| **Average** | **574 ± 344** | **660 ± 376** | **629 ± 391** | **-** |

### Comparison of pulsed conditions

The normalized MEP amplitudes for each parameter set are shown in **Figure 5A** ordered from greatest inhibition to greatest excitation. Parameter set 2 (6 W/cm^2^ I_SPPA_, 10% DC, 100 msec) resulted in the greatest excitation (110% ± 11.8%). Parameter set 8 (6W/cm^2^ I_SPPA_, 30% DC, 500 msec) resulted in the greatest inhibition (70% ± 0.28%). To test for statistical significance for which parameters were significantly different from baseline we tested each set using a one-sample t-test Bonferroni corrected for multiple comparisons (0.05/20) = 0.0025. Only sets 8, 16 and 10 met this rigid statistical threshold (see **Figure 5A**). Set 8: t(17) = 4.44, p = 3.6 *10e-4, η^2^ = 0.54; Set 16: t(17) = 4.52, p = 3.02*10e-4, η^2^ = 0.53; Set 10: t(17) = 3.60, p = 0.0022, η^2^ = 0.66. The mean ± SD for condition 16 (24 W/cm^2^; 1% DC; 500 msec) was 0.75% ± 0.23%. The mean ± SD for condition 10 was 0.75% ± 0.29%. Probability histograms for each of these conditions as well as the next condition that showed the greatest inhibition (Condition 17: t(17) = −2.49, p = 0.023) that did not meet statistical significance are presented in **Figure 5B & Table 3**.

**Figure 5.**
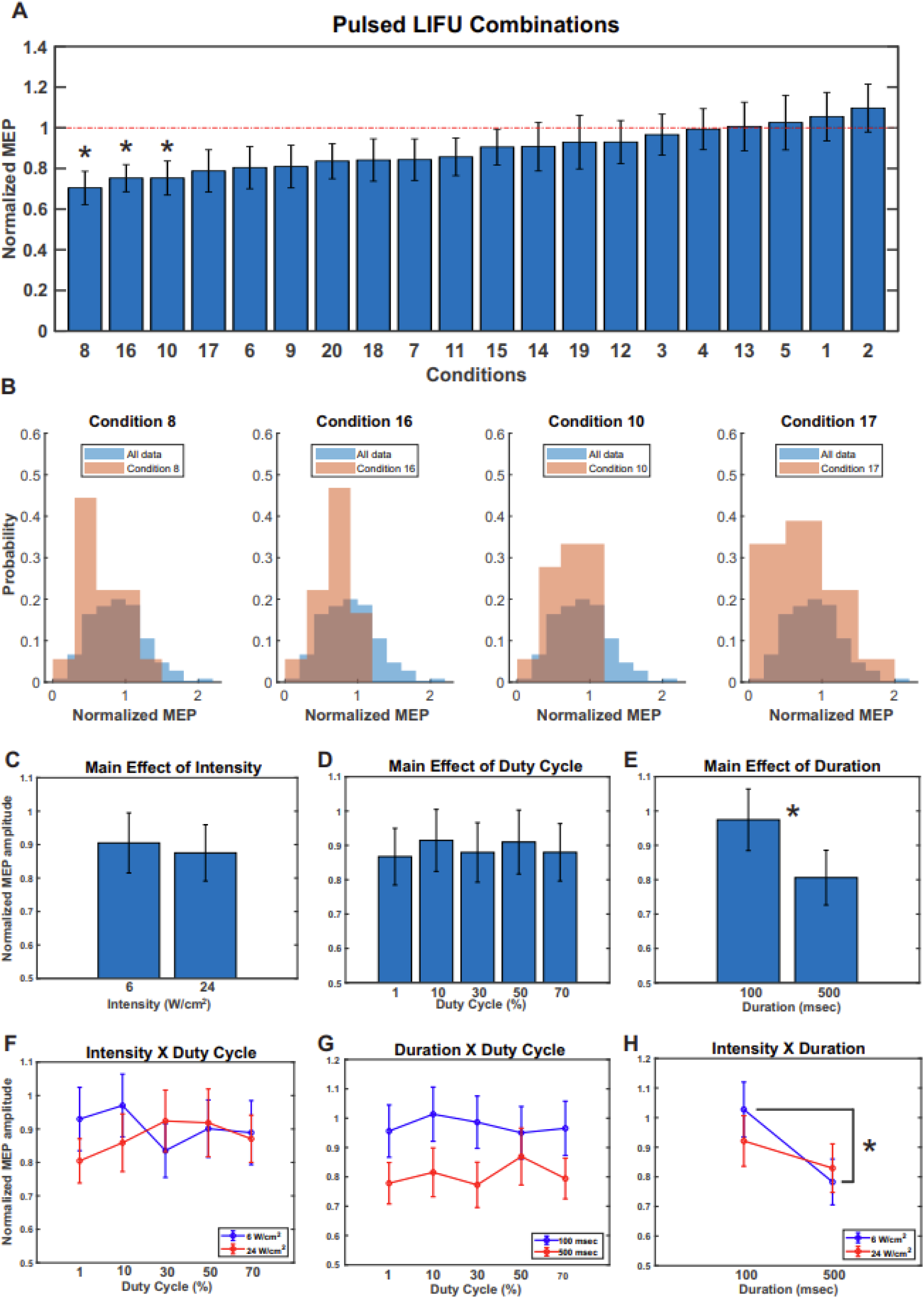
Pulsed parameter combinations. **A.** Group (N = 18) mean ± SEM bar plot depicting normalized MEP amplitudes for all 20 pulsed combinations of ultrasound parameters, sorted from inhibition to excitation. The parameters associated with each condition number shown in **Table 3**. Asterisks denote a significant difference from baseline at p < 0.0025. **B.** Probability histograms for the three conditions shown to significantly differ from baseline (8, 16, 10) and the next condition that showed the greatest inhibition but did not meet statistical significance (17). **C.** Main effect of Intensity. **D**. Main effect of duty cycle. **E**. Main effect of duration. Asterisk denotes a significant difference between groups at p < 0.0001. **F**. 2-way interaction between intensity and duty cycle. **G**. 2-way interaction between duration and duty cycle. **H**. 2-way interaction between intensity and duty cycle. Asterisk denotes a significant difference between groups at p < 0.05.

**Table 3:**
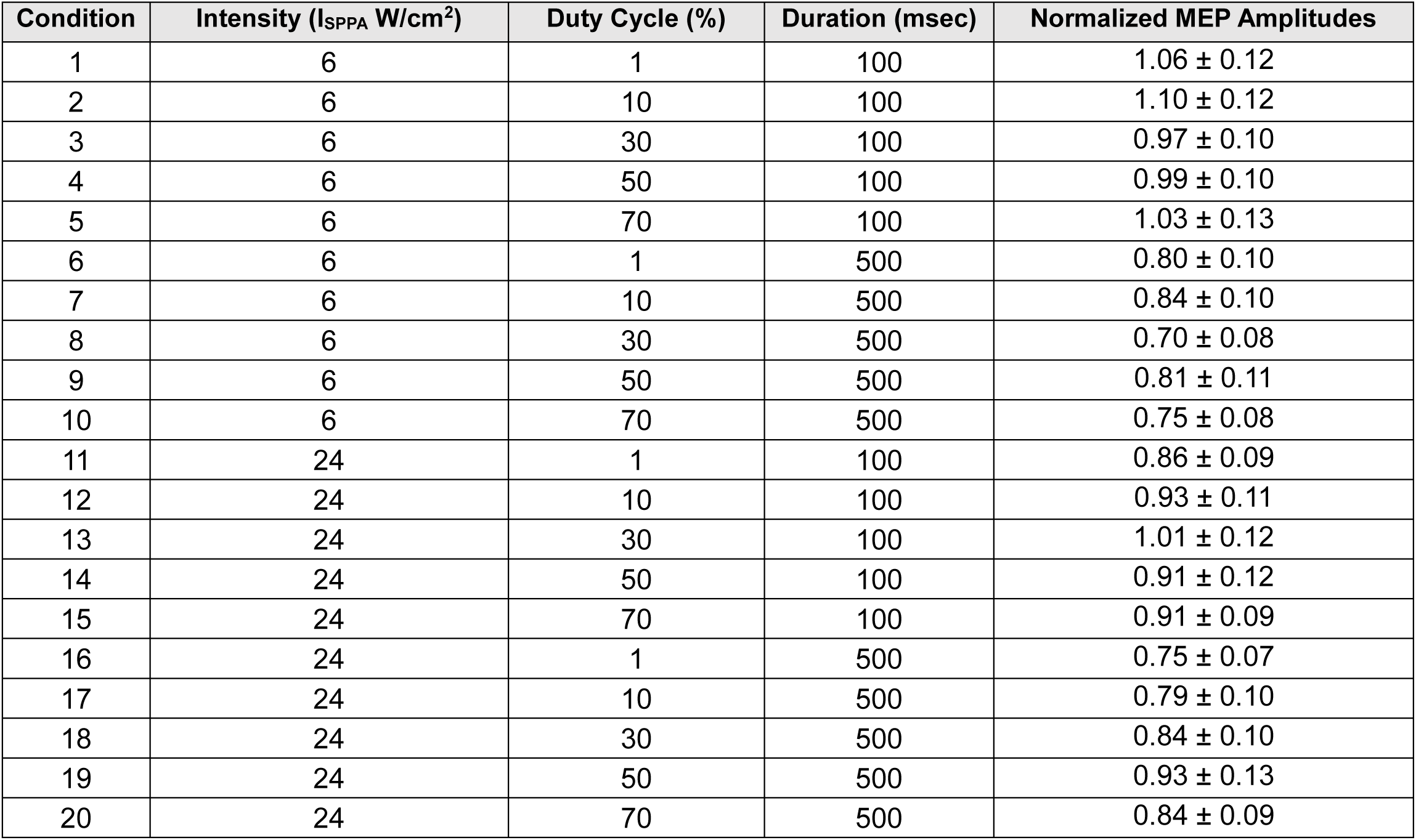
Pulsed Parameter Combinations: parameter values and corresponding mean±SD normalized MEP amplitudes.

| Condition | Intensity (I<sub>SPPA</sub> W/cm<sup>2</sup>) | Duty Cycle (%) | Duration (msec) | Normalized MEP Amplitudes |
| --- | --- | --- | --- | --- |
| 1 | 6 | 1 | 100 | 1.06 ± 0.12 |
| 2 | 6 | 10 | 100 | 1.10 ± 0.12 |
| 3 | 6 | 30 | 100 | 0.97 ± 0.10 |
| 4 | 6 | 50 | 100 | 0.99 ± 0.10 |
| 5 | 6 | 70 | 100 | 1.03 ± 0.13 |
| 6 | 6 | 1 | 500 | 0.80 ± 0.10 |
| 7 | 6 | 10 | 500 | 0.84 ± 0.10 |
| 8 | 6 | 30 | 500 | 0.70 ± 0.08 |
| 9 | 6 | 50 | 500 | 0.81 ± 0.11 |
| 10 | 6 | 70 | 500 | 0.75 ± 0.08 |
| 11 | 24 | 1 | 100 | 0.86 ± 0.09 |
| 12 | 24 | 10 | 100 | 0.93 ± 0.11 |
| 13 | 24 | 30 | 100 | 1.01 ± 0.12 |
| 14 | 24 | 50 | 100 | 0.91 ± 0.12 |
| 15 | 24 | 70 | 100 | 0.91 ± 0.09 |
| 16 | 24 | 1 | 500 | 0.75 ± 0.07 |
| 17 | 24 | 10 | 500 | 0.79 ± 0.10 |
| 18 | 24 | 30 | 500 | 0.84 ± 0.10 |
| 19 | 24 | 50 | 500 | 0.93 ± 0.13 |
| 20 | 24 | 70 | 500 | 0.84 ± 0.09 |

We next examined how and if parameters interacted to produce effects. The three-way repeated measures ANOVA revealed no significant main effect of INTENSITY (F(1, 358) = 0.61, p = 0.44, η_p_ ^2^ = 0.0018) (**Fig. 5C**) or DUTY CYCLE (F(4, 356) = 0.24, p = 0.92, η_p_ ^2^ = 0.0028) (**Fig. 5D**) and a significant main effect of DURATION (F(1, 358) = 19.36, p < 0.0001, η_p_ ^2^ = 0.0539) (**Fig. 5E**). There was no significant interaction between INTENSITY x DUTY CYCLE (F(4, 356) = 1.1, p = 0.3577, η_p_ ^2^ = 0.0127) (**Fig. 5F**), or DUTY CYCLE x DURATION (F(4, 356) = 0.36, p = 0.83, η_p_ ^2^ = 0.0043) (**Fig. 5G**). There was an interaction of DURATION x INTENSITY (F(1, 358) = 4.0, p = 0.0463, η ^2^ = = 0.0116) (**Fig. 5H**). The three-way interaction INTENSITY x DUTY CYCLE x DURATION was not significance (F(4, 356) = 1.1, p = 0.35, η_p_ ^2^ = 0.0010). Post-hoc testing of the interaction between intensity and duration revealed significant differences between the low-intensity/short-duration condition and both the low-intensity/long-duration (p < 0.0001) and high-intensity/long-duration conditions (p = 0.0014), with the high-intensity/short-duration condition trending towards significance (p = 0.0511). These results show that for short durations, lower intensity produces less inhibition compared to higher intensity but at longer durations, the lower intensity condition becomes more inhibitory than the higher intensity condition (**Fig. 5H**).

### Pulsed vs. Continuous

The second phase of the study evaluated four continuous-wave conditions with two different durations (70 msec and 150 msec) and two intensity levels (I_SPPA_: 6 W/cm² and 24 W/cm²). These were compared to cycle-matched pulsed LIFU conditions at both I_SPPA_ levels: one with a 70% duty cycle for 100 msec (set 5 & 15), and another with a 30% duty cycle for 500 msec (set 8 & 18). The cycle-matched pulsed and continuous condition pairs are as follows: 5 with 21, 15 with 22, 8 with 23, and 18 with 24 (refer to **Fig. 6A** and **Table 4**). The mean ± SD of normalized MEP amplitudes for all subjects across the 4 continuous and 4 pulsed conditions was 0.87 ± 0.13, ranging from 0.70 to 1.03 and 0.87 ± 0.13 and 1.00 ± 0.11, ranging from 0.84 to 1.10, respectively (**Fig. 6A**) and parameter specific responses are listed in **Table 4**. The 3-way repeated measures ANOVA revealed a three-way interaction between INTENSITY x PULSING x NUMBER OF CYCLES (F(1, 358) = 5.01, p = 0.027, η_p_ ^2^ = 0.0378. Post-hoc testing of the interaction revealed significant differences between the pulsed long-duration low-intensity and continuous wave long-duration low-intensity conditions (p = 0.024). This analysis indicates that the inhibitory effect of the pulsing scheme depends on both the duration and intensity of the treatment. Specifically, pulsed LIFU is more inhibitory than CW LIFU only at longer durations and lower intensities. At shorter durations, there is no significant difference between the two schemes at either intensity (**Figure 6H**). The main effects and interactions of each factor are present in **Figure 6 B-G**. Because there was a 3-way interaction these effects were not explored but results of the omnibus ANOVA for each factor main effect and interaction are below: There were no significant main effect of INTENSITY (F(1, 358) = 0.53, p < 0.47) (**Fig. 6B**) or NUMBER OF CYCLES (F(1, 358) = 0.15, p = 0.70) (**Fig. 6D**). However, there was a significant main effect of PULSING (F(1, 358) = 5.33, p < 0.023) (**Fig. 6C**) as well as a significant interaction between PULSING x NUMBER OF CYCLES (F(1, 358) = 4.05, p = 0.046) (**Fig. 6G**). Further analysis revealed no significant interactions between INTENSITY x PULSING (F(1, 358) = 0, p = 0.99), or INTENSITY x NUMBER OF CYCLES (F(1, 358) = 0.06, p = 0.80) (**Fig. 6F**).

**Figure 6.**
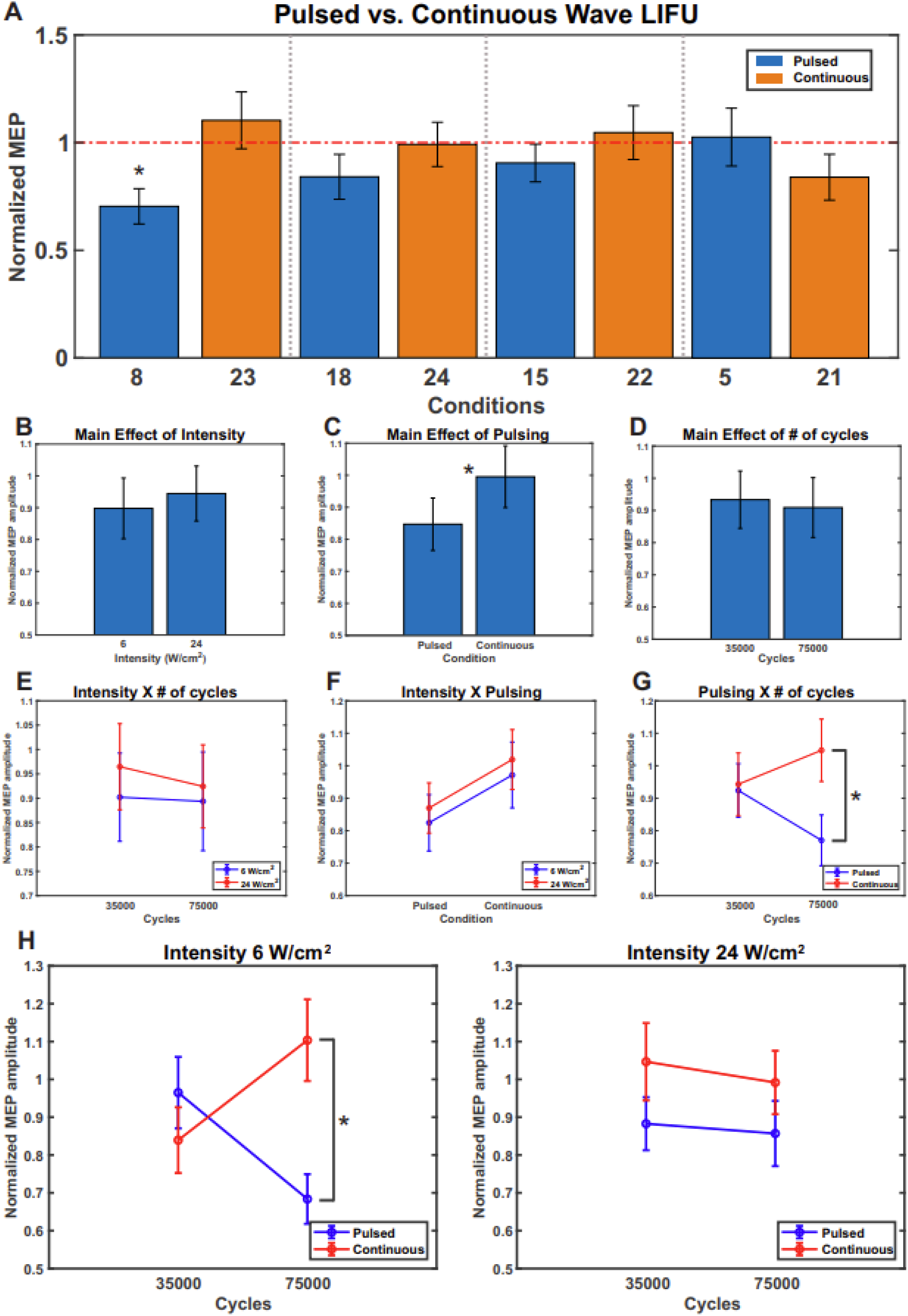
Pulsed vs. Continuous. **A.** Group (N = 17) mean ± SEM bar plot depicting normalized MEP amplitudes for all 8 pulsed vs. continuous conditions, grouped into cycle-matched pairs between the two pulsing schemes. The parameters associated with each condition number shown in **Table 4**. **B.** Main effect of intensity. **C**. Main effect of pulsing. Asterisk denotes a significant difference between groups at p < 0.05. **D**. Main effect of number of cycles. **E**. 2-way interaction between intensity and number of cycles. **F**. 2-way interaction between intensity and pulsing. **G**. 2-way interaction between pulsing and number of cycles. Asterisk denotes a significant difference between groups at p < 0.05. **H.** 3-way interaction between intensity, pulsing, and number of cycles for the lower **and** higher intensity condition (left and right, respectively). Asterisk denotes a significant difference between groups at p < 0.05.

**Table 4:** Pulsed vs. Continuous parameter combinations: parameter values and corresponding mean±SD normalized MEP amplitudes.

| Intensity | # Cycles | Pulsed Condition | Normalized MEP Amplitudes | CW Condition | Normalized MEP Amplitudes |
| --- | --- | --- | --- | --- | --- |
| 6 | 75000 | 8 | 0.70 ± 0.08 | 23 | 1.10 ± 0.13 |
| 24 | 75000 | 18 | 0.84 ± 0.10 | 24 | 0.99 ± 0.10 |
| 24 | 35000 | 15 | 0.91 ± 0.09 | 22 | 1.05 ± 0.13 |
| 6 | 35000 | 5 | 1.03 ± 0.13 | 21 | 0.84 ± 0.11 |

### Safety

A review of symptoms questionnaire [14] queried the participants on the presence of various symptoms and their severity (absent / mild / moderate / severe) scored on a scale of 1-4. This questionnaire was collected at the beginning of each visit which had LIFU/TMS application and 30 minutes after TMS/LIFU application to indicate any changes of from the intervention. **Figure 7** depicts all sessions in which participants received TMS or LIFU/TMS, even if they had a failed session. If there was no change, or improvement in the participant’s symptoms, the symptoms were represented as absent. There were a few sessions in which participants had mild neck pain, sleepiness, headache, and scalp sensations. Of the 16 participants who responded to the one-month follow-up questionnaire, none of them reported any persisting or new symptoms.

**Figure 7.**
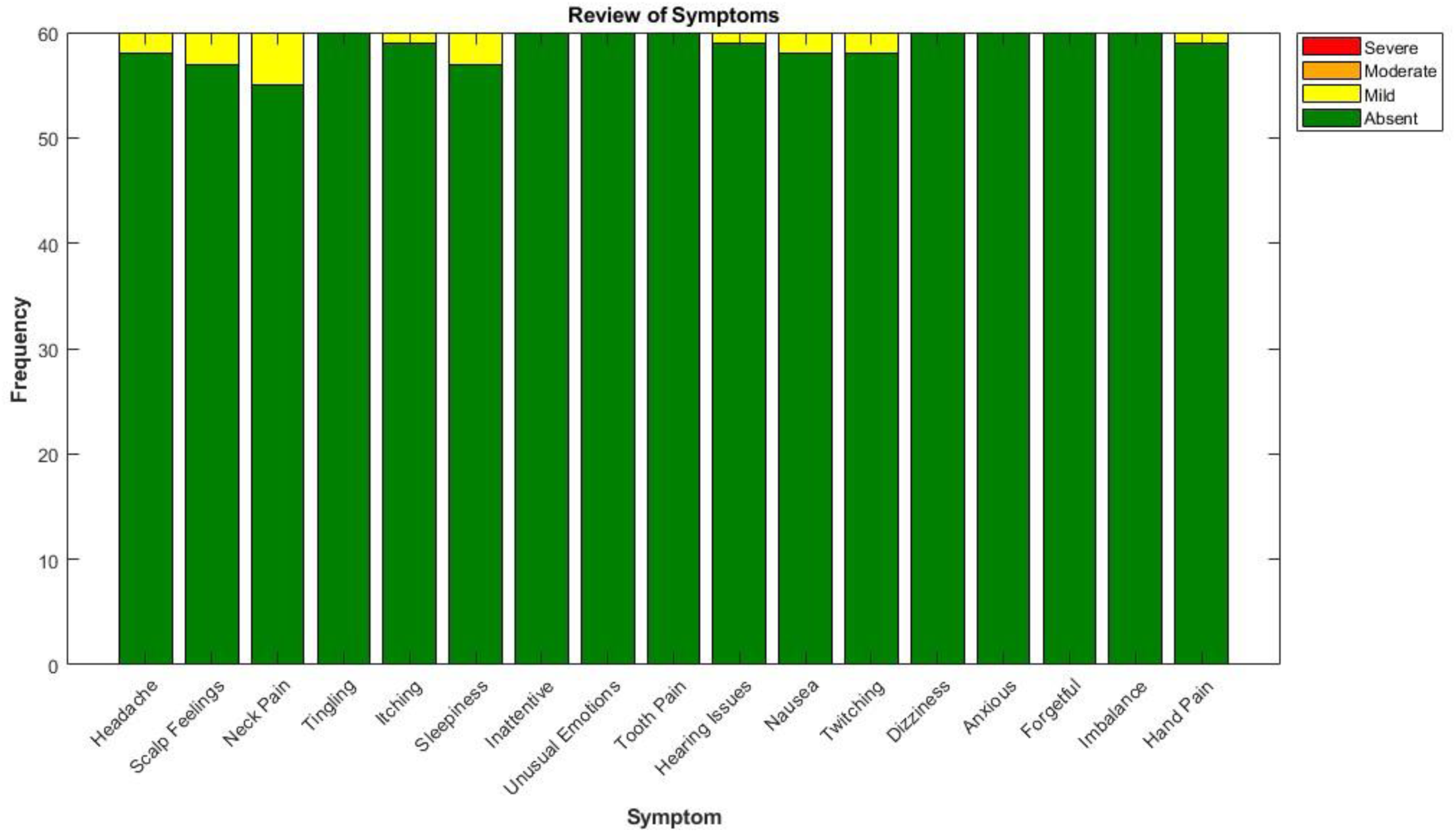
Report of Symptoms questionnaire results. Questionnaire was collected before and after any session involving TMS or LIFU/TMS stimulation. Each bar represents the total count of symptoms across all participant sessions (60 sessions) and the colors illustrate symptom severity. If there was no change or improvement in the participant’s symptoms, the symptoms were represented as absent (green).

## DISCUSSION

This study explored how different LIFU parameters interact to affect motor cortex excitability assessed by the amplitude of the TMS derived motor evoked potential recorded from the FDI muscle. We first examined 20 different LIFU parameter combinations delivered in pulse mode at a PRF of 1kHz. We followed this up by comparing the effects of 4 selected pulsed conditions with 4 cycle-matched continuous-wave LIFU parameter sets. We found that some parameter sets induced inhibition whereas others showed no significant effect compared to baseline. None of the parameter sets resulted in an increase in amplitude of the MEP. Furthermore, for pulsed LIFU, the degree of inhibition was influenced by both duration and intensity, with lower intensity LIFU being more inhibitory than higher intensity LIFU at longer durations. When comparing pulsed and continuous wave LIFU, the results demonstrated that the magnitude of the effect varied depending on the pulsing scheme, duration, and intensity. Specifically, the largest effects were the result of pulsed LIFU at lower intensities and longer durations.

### Pulsed Parameter Combinations

We originally hypothesized that lower duty cycles (1 – 30%) would have inhibitory effects, while higher duty cycles (50 & 70%) would produce excitatory effects based on prior literature [34]. We further expected that increasing the total energy delivered—through longer durations or higher intensities—would amplify any effects under the hypothesis that more total energy would serve to amplify duty cycle effects. However, our results did not find any conditions to produce significant excitation from the baseline. Instead, parameter combinations resulted in either inhibition or no effect. The condition achieving the greatest inhibition was produced by the lowest intensity (I_SPPA_ 6W/cm^2^) and longest duration (500 msec) at an intermediate duty cycle (30%). Notably, varying the duty cycle was not found to significantly modulate the magnitude of neuromodulation across conditions. These findings contradict our hypothesis that the magnitude of the LIFU effect would be dependent on the duty cycle due to increased energy deposition. Moreover, the observation that lower intensity LIFU produced a greater inhibitory effect than higher intensity LIFU over longer durations suggests that total energy delivered may not be the key factor in determining the degree of effect. Instead, the degree of neuromodulation may depend on both the duration of the intervention and the intensity level.

Previous studies examining the LIFU parameter space in humans and large animals have observed duration-dependent effects of LIFU on MEP amplitudes with varying influences of intensity and duty cycle, appearing to depend on the experimental design. When delivered concurrently in human subjects, durations of 400 and 500 milliseconds both resulted in a significant reduction of MEP amplitudes compared to the sham control, whereas shorter durations failed to produce consistent inhibition [13]. In this study, when interleaving the parameter sets of interest, only a duty cycle of 10% (0.5s duration, 1000Hz PRF, I_SPPA_ = 2.32 W/cm^2^) resulted in significant inhibition from sham and the effect of varying intensity was not explored. Another study targeting the M1 of anesthetized sheep with LIFU demonstrated a more effectively reduced muscle twitch response rates at lower intensities compared to higher intensities (I_SPPA_: 15.8, 18.2 W/cm^2^) which matched up with our findings [31]. However, contrary to our results, this study also found that the shortest duration condition (0.5 sec) produced the greatest effect compared to longer durations (2 and 3 seconds).

A second human study examining the LIFU parameter space that both extending the offline ultrasound delivery period and increasing the duty cycle enhanced motor cortical excitability for up to 90 minutes, indicating that the degree of neuromodulation may depend on the total energy delivered [32]. While a longer sonication duration led to a greater enhancement in cortical excitability, there was no significant difference in MEP amplitude between the 10% and 15% duty cycles, suggesting a non-linear relationship between duty cycle and effect. Additionally, when the intensity was reduced from 20 W/cm^2^ to 10 W/cm^2^ I_SPPA_, motor cortical excitability was no longer significantly changed from baseline. This discrepancy between these results and our study may be due to the PRF used during that portion of the study (5 Hz) which is thought to facilitate MEPs; however, independent attempts to replicate these findings have been unsuccessful in generating excitation [44] and further research is needed to determine whether LIFU can effectively facilitate brain activity.

### Effect of PRF

Throughout the study, we used a constant PRF of 1kHz based on our previous work and on PRFs used in other LIFU studies [10,13,18,33]. We were additionally guided by theoretical models which were developed to help explain results found in small animal models, suggesting that PRF does not play a major role in determining the overall effects of LIFU. Previous research across various models—including rodent brain slices, single cells, small animals, and human subjects—has demonstrated that varying PRF affects the magnitude of neuromodulation [3,27,28,33]. In the murine motor cortex, a 1500 Hz PRFs condition produced a greater change in neural response rates compared to the lower 300 Hz PRFs [27]. In single-cell studies, LIFU may differentially target inhibitory or excitatory neuronal subpopulations depending on the PRF value chosen [28]. In rodents, higher PRFs increase the EMG failure rate when stimulating paw twitches with LIFU [3]. Similarly, in human subjects, PRF-dependent effects on MEP inhibition have been observed where 100 Hz is most effective, 10 Hz is moderately effective, and 1000 Hz had no effect compared to a sham treatment [33]. This last observation is intriguing given that our stimulation protocol successfully utilized 1000 Hz PRF to induce MEP amplitude inhibition; however, this comparison is limited since Zeng et al. 2024 was conducted using an offline LIFU protocol and the present study was conducted in an online design [32]. Therefore, it remains unclear how the findings related to duration and the interaction between duration and intensity might be influenced when PRF is varied. Moreover, it is possible that parameters that are unresponsive in the current study might exhibit different outcomes under varying PRF conditions. This gap underscores the importance of protocol differences and their potential impact on study outcomes. It was a missed opportunity in this paper to test levels of PRF and how they interact with the other parameters tested.

### Pulsed vs. Continuous

We hypothesized that pulsed and continuous LIFU would confer the same degree of effect and that, for both of these conditions, only the total amount of energy would define the degree of neuromodulation. Our results, however, demonstrate that the overall magnitude of the effect is contingent upon how long, at what intensity, and under what scheme (pulsed or continuous) the ultrasound was delivered. Specifically, a significant difference was found between the long-duration low-intensity pulsed and long-duration low-intensity continuous wave conditions of ∼40%. This finding suggests that different stimulation modes, particularly between pulsed and continuous waveforms, may produce varying effects, especially in situations characterized by lower intensity and prolonged duration. Previous studies have explored the efficacy of continuous versus pulsed LIFU for neuromodulation in *ex vivo* samples and small animals. In salamander retinas, continuous wave LIFU was found to be more effective for modulating retinal cells compared to pulsed LIFU [35]. Similarly, when targeting the motor cortex in anesthetized mice to induce paw twitches, continuous wave LIFU was more effective at generating paw twitches than pulsed LIFU at higher intensities [4]. However, our study presents contrasting findings. We observed that pulsing was more effective at inhibiting MEP amplitudes for the longer duration, lower intensity LIFU. These differences may arise from variations across the model systems and warrants further investigation to determine the role of pulsing schemes in the overall degree of neuromodulation.

### Safety

Mild symptoms such as neck pain, sleepiness, headache, and scalp sensations were reported though the overall frequency and severity of symptoms were low. Neck pain and sleepiness experienced by the participants may be attributed to the requirement of sitting still and keeping their heads in a fixed position during the study visit. These symptoms are not surprising and match well with the symptoms previously reported in ultrasound research [14]. Importantly, during the one-month follow-up, none of the 16 participants who responded reported any new or persistent symptoms. This absence of long-term adverse effects suggests a favorable safety profile for the LIFU/TMS interventions used in this study.

### Heating

While previous research has explored the role of thermal effects on neuroinhibition [45], the results of the present study do not seem to be influenced by temperature changes. Darrow et al. (2019) found that among all the tested LIFU parameters and experimental conditions, the increase in brain temperature caused by ultrasound was the strongest predictor of neural inhibition, with significant effects observed at temperatures above 0.5 °C. However, these findings do not apply to our study for two key reasons. First, the most conservative estimate of thermal rise in the brain during our study remained below 0.5 °C. Second, the parameter sets most effective at inhibiting motor cortical excitability were not the ones with higher intensity and duty cycle, but rather those with lower intensity or lower duty cycle.

### Limitations

This study has several limitations. The chosen parameter sets were limited due to either potential risks due to thermal bioeffects (i.e. longer durations or higher intensities) or overall study time constraints. For example, participants requiring higher TMS output (i.e., 100%) to achieve the MEP threshold limited the number of settings we could test per day, as cooldown periods were necessary – thus different subjects received different total number of tested parameters within session but there was no systematic effect and this likely did not contribute to findings. Furthermore, new baseline data was taken for each session also helping to mitigate this potential. There was the potential for carry-over effects with our study design. Previous research used short online LIFU and demonstrated no cumulative effects [11,46]. Nevertheless, we took precaution to make sure there were no carry-over effects between parameter sets by using go/no-go MEP amplitude criteria. In some cases, extended inhibition was found.

Additionally, LIFU has been shown to produce auditory effects that may interfere with its neuromodulatory impacts [47–50]. We did not employ an auditory mask in these experiments as each active condition would serve as a matched auditory control. We are confident that the results are not due to non-specific auditory confounds, as higher amplitudes and longer durations—conditions known to produce the most significant auditory artifacts [51]—are typically associated with greater cortical excitation. If auditory artifacts were influencing our findings, we would expect the greatest inhibition to occur under these conditions. However, this pattern is not observed in our results.

While our focus was primarily on comparing different parameter sets, incorporating an inactive sham condition to demonstrate no effects could also have strengthened our comparisons. To mitigate the lack of an inactive sham condition, sessions were randomly ordered to reduce potential biases in data collection and analysis. Another potential limitation was the variation in individual subject attenuation of the ultrasound energy which may contribute to differences in effects of the same condition across subjects, thus making interpretation of the comparison between ‘low’ and ‘high’ LIFU challenging. Nevertheless, since pressure attenuation at the skull bone is linear for each participant, low/high distinctions within-subject remain valid. Lastly, the study is limited by the range of pressures evaluated, which averaged between 141 and 275 kPa. Several promising LIFU studies have been conducted at higher pressures, near 1 MPa [21,22,24]. As the field advances toward clinical applications of LIFU, it will be important to characterize the parameter space at these pressures which approach 1 MPa.

## CONCLUSIONS & FUTURE WORK

LIFU to M1 in healthy humans, regardless of parameter combinations in the range employed, either conferred inhibition or had no significant effect. Significant excitation was not observed. In general, longer durations and lower intensities looks to be more efficacious for inhibition independent of duty cycle when delivered at a 1 kHz PRF. In addition, pulsed LIFU was more efficacious for inhibition as compared to continuous LIFU, also depending on the duration and intensity of the sonication. Finally, it is not clear how and if ‘online’ parameters transfer to so-called ‘offline’ protocols that may induce greater or lasting neuromodulatory effects and warrants future study.

