## Supplemental Tables for "Examination of low-intensity focused ultrasound parameters for human neuromodulation"

**Table 1**: Parameter conditions completed per study visit.

| Participant # | Parameters Completed | | | |
|---|---|---|---|---|
| | *Visit 1* | *Visit 2* | *Visit 3* | *Visit 4* |
| 1 | 7 | 10 | 0 | 7 |
| 2 | 2 | 9 | 9 | 4 |
| 3 | 2 | 5 | 13 | |
| 4 | 3 | 13 | 8 | |
| 5 | 10 | 8 | 6 | |
| 6 | 11 | 9 | 4 | |
| 7 | 7 | 17 | | |
| 8 | 12 | 12 | | |
| 9 | 12 | 12 | | |
| 10 | 12 | 12 | | |
| 11 | 12 | 12 | | |
| 12 | 12 | 12 | | |
| 13 | 12 | 12 | | |
| 14 | 12 | 12 | | |
| 15 | 13 | 11 | | |
| 16 | 13 | 11 | | |
| 17 | 14 | 10 | | |
| 18 | 15 | 9 | | |

**Table 2**: Go/no go protocol enactment by parameter set across visits.

| Participant # | Visit | date | Parameter Tested | | | | | | |
|---|---|---|---|---|---|---|---|---|---|
| | | | 1st | 2nd | 3rd | 4th | 5th | 6th | 7th |
| 1 | visit 1 | 3/14/2023 | 18 | - | 6 | - | 7 | 21 | 17 |
| 2 | visit 2 | 3/15/2023 | 19 | - | 17 | - | - | - | - |
| 3 | visit 1 | 3/17/2023 | 7 | - | - | - | - | - | - |
| 4 | visit 1 | 3/10/2023 | - | 8 | - | - | - | 16 | 9 |
| 5 | visit 2 | 4/5/2023 | - | - | - | 18 | - | - | - |
| 6 | visit 1 | 3/7/2023 | - | - | - | - | - | - | 19 |
